# How Much Does Electrode Density Matter? A Multilevel Evaluation from EEG Signals to Cortical Networks

**DOI:** 10.64898/2026.09.26.754643

**Authors:** Aya Kabbara, Marc Vérin, Mahmoud Hassan

**Affiliations:** Graduate Studies and Research, Lebanese American University, Beirut, Lebanon; Department of Computer Science, Lebanese International University, Tripoli, Lebanon; Brain-Clinical and Experimental Neuroplasticity (B-CLINE), Laboratoire Interdisciplinaire pour l’Innovation et la Recherche en Santé d’Orléans (LI2RSO), Université d’Orléans, Orléans, France; Neurology Department, CHU d’Orléans, Orléans, France; MINDIG, Rennes, France; Institute of Biomedical and Neural Engineering, School of Science and Engineering, Reykjavik University, Reykjavik, Iceland

**Keywords:** high-density EEG, electrode density, spatial sampling, EEG microstates, effective connectivity, traveling waves, source reconstruction, brain networks

## Abstract

**Background:** High-density electroencephalography (EEG) improves spatial sampling but entails greater acquisition cost, preparation time, participant burden, and computational demand. However, the extent to which electrode reduction affects successive levels of EEG analysis from global signal properties to cortical network inference remains insufficiently understood.

**Methods:** We systematically subsampled two independent resting-state EEG datasets comprising 120 children recorded with 128 channels and 98 adults recorded with 256 channels. Density-dependent effects were assessed across 128-, 256-, 64-, 32-, and 19-channel configurations using a multilevel framework encompassing signal representation and quality, global and regional spectral power, EEG microstates, scalp-level effective connectivity, traveling waves, and source-level cortical hub topology.

**Results:** Sensitivity to electrode reduction increased with the spatial specificity of the outcome. Global relative spectral power and explained variance were comparatively stable across configurations. In contrast, signal-to-noise ratio, entropy, reconstruction error, regional spectral distributions, traveling-wave estimates, effective connectivity, and microstate dynamics became progressively more density-sensitive. Although canonical microstate classes remained broadly identifiable at lower densities, their global explained variance, temporal parameters, and transition structure were altered. Source-level hub identity and centrality ranking showed the greatest vulnerability, with markedly reduced cross-density agreement at 32 and 19 channels. Across both datasets, several spatially dependent measures showed a transition around 64 channels, whereas estimates obtained with 128 and 256 channels were generally more similar.

**Conclusions:** Electrode density affects EEG outcomes according to the spatial scale of the intended inference. Sparse montages may preserve broad global summaries but are less reliable for regional, spatiotemporal, and source-network analyses. Under the resting-state conditions examined here, approximately 64 channels represented a pragmatic transition for several spatially dependent measures. This value should be interpreted as context-specific guidance rather than a universal threshold.

**Highlights:**

- Electrode-density effects increase with the spatial specificity of the EEG measure.
- Global spectral power and explained variance remain comparatively robust at low density.
- Microstate classes persist at low density, but temporal dynamics and syntax are distorted.
- Connectivity, traveling waves, and cortical hub inference are more sensitive to sparse sampling.
- Across two independent datasets, many spatially dependent measures deteriorate below approximately 64 channels.

## Introduction

Electroencephalography (EEG) has been a cornerstone of neuroscientific research and clinical practice since Hans Berger first recorded human brain electrical activity nearly a century ago (Berger, 1929). From early recordings obtained with only a few scalp electrodes, EEG has evolved into sophisticated multichannel systems capable of sampling electrical brain activity from hundreds of locations simultaneously. Alongside complementary neuroimaging modalities such as magnetic resonance imaging and positron emission tomography, EEG provides a distinctive combination of millisecond temporal resolution, non-invasive acquisition, and direct sensitivity to neuronal electrical activity, making it particularly well suited to the study of rapidly evolving brain dynamics (Niedermeyer & da Silva, 2005; Logothetis, 2008). Advances in acquisition hardware, signal processing, and computational modelling have further extended EEG analysis beyond conventional waveform inspection toward spatial mapping, source reconstruction, functional and effective connectivity, and network-level characterization (Niedermeyer & da Silva, 2005; Michel & Murray, 2012, Hassan et al., 2018).

A central development in this evolution has been the advent of high-density EEG (hd-EEG), defined by configurations exceeding 128 channels (Lindsey et al., 2017). By sampling the scalp electric field at a larger number of locations, dense electrode arrays can provide a more complete representation of its spatial structure and improve the conditions under which cortical generators are estimated. In epilepsy, for example, hd-EEG improves the characterization and localization of epileptiform activity and has enabled the reconstruction of distributed interictal epileptic networks from non-invasive recordings (Yamazaki et al., 2013; Hassan et al., 2017; Danoun et al., 2019; Stoyell et al., 2021). More generally, adequate spatial sampling is an important determinant of EEG source-localization accuracy (Michel & He, 2011; Song et al., 2015), and dense-array recordings have supported the development of source-connectivity approaches aimed at reconstructing large-scale cortical networks from scalp EEG (Hassan et al., 2014; Allouch et al., 2022, Allouch et al., 2023).

The advantages of increasing electrode density, however, come at a practical cost. Dense systems require more expensive equipment, longer preparation and acquisition times, greater participant tolerance, and increased computational and storage resources (Michel et al., 2004). Consequently, conventional 19- or 32-channel systems remain common in clinical and research environments, while 64-, 128-, and 256-channel configurations are used when greater spatial resolution is required. This creates a fundamental methodological question: how much spatial sampling is actually required to preserve a given EEG result? Importantly, the answer may depend on what is subsequently inferred from EEG signals.

Several studies have addressed specific aspects of this question, providing evidence that the consequences of electrode reduction are measure-dependent. At the scalp level, theoretical, simulation, and empirical work has shown that electrode density influences spatial sampling accuracy and can affect measured EEG properties, including spectral estimates (Grieve et al., 2004; Greischar et al., 2004). For spatially organized EEG features, microstate analyses have shown comparatively stable solutions at moderate densities but reduced reliability with sparse electrode configurations (Zhang et al., 2021). At the source level, electrode density is a recognized determinant of source-localization accuracy (Lantz et al., 2003; Song et al., 2015). Simulation-based investigations have further demonstrated that channel density interacts with the inverse solution and connectivity estimator in determining the accuracy of reconstructed EEG source-space and resting-state networks (Hassan et al., 2017, Allouch et al. 2022, 2023).

Empirical evidence at the network level is consistent with these observations. Hatlestad-Hall et al. (2023), using resting EEG from 33 participants recorded with 128 electrodes and subsequently reduced to 64, 32, and 19 channels, showed that source-level functional connectivity and graph-theoretical measures became progressively less consistent with the high-density solution as electrode number decreased. Their results suggested that approximately 64 electrodes provided a reasonable compromise for source-level graph analysis under the conditions examined. More recently, Dattola et al. (2026) compared 173-, 64-, and 18-channel configurations in resting-state recordings from 17 individuals with mild cognitive impairment and found that the high-density configuration provided the clearest network-level relationships with cognitive performance, further suggesting that electrode density can affect the sensitivity of source-level network analyses.

Together, these studies establish that electrode density can affect EEG analysis, but they also reveal an important limitation of the existing literature. Previous investigations have generally focused on individual stages or specific families of measures: spatial sampling error (Grieve et al., 2004), scalp spectral power (Greischar et al., 2004), microstate reliability (Zhang et al., 2021), source localization (Lantz et al., 2003), or source-level functional connectivity (Hassan et al., 2017, Allouch et al., 2022, 2023) and graph-theoretical properties (Hatlestad-Hall et al., 2023). Moreover, several influential methodological studies have relied on simulations, whereas empirical investigations have typically involved relatively small cohorts or focused on a specific clinical or analytical endpoint. Consequently, it remains unclear whether there is a single electrode-density requirement for resting-state EEG or whether sensitivity to spatial undersampling increases systematically as the analysis progresses from global signal properties toward increasingly spatially specific physiological and cortical-network inference.

Here, we address this question using two independent, comparatively large empirical resting-state EEG datasets (N= 218) and a multilevel analytical framework. Rather than evaluating a single EEG endpoint, we examine electrode-density effects across a hierarchical analytical framework. At the first level, we quantify changes in signal representation and quality using signal-to-noise ratio, entropy, explained variance, and reconstruction error. At the second level, we examine complementary physiological properties directly from scalp EEG, encompassing oscillatory content through absolute and relative spectral power, recurrent global field configurations through EEG microstates, directed multivariate interactions through MVAR/DTF, and spatiotemporal propagation through traveling-wave analysis. These measures were intentionally evaluated at the sensor level to characterize increasingly spatial physiological information without introducing the additional assumptions associated with solving the EEG inverse problem. At the third level, we introduce source reconstruction and assess the most spatially specific inference considered here: whether electrode density alters cortical functional connectivity and the identity and consistency of network hubs. .

By applying this multilevel framework to two comparatively large, independent empirical datasets, we test the hypothesis that the consequences of electrode reduction are not uniform across EEG measures but increase with the spatial specificity of the inference being made. Our objective is therefore to determine which properties of resting-state EEG remain robust under spatial undersampling and which require denser spatial sampling. Such evidence may help guide the selection of EEG systems according to the intended analytical endpoint, balancing physiological and anatomical fidelity against acquisition complexity, cost, and accessibility.

## Methods

To evaluate the effect of spatial sampling on EEG data quality, feature extraction, and network-level inference, we analyzed two independent resting-state EEG datasets encompassing 218 participants. Each dataset was processed across multiple electrode density configurations using identical preprocessing and analysis pipelines, allowing within-subject comparisons that are free of inter-subject confounds. The analytical framework was organized into three hierarchical levels: signal quality, physiological features, and cortical network topology. The full pipeline is illustrated in Figure 1.

**Figure 1.**
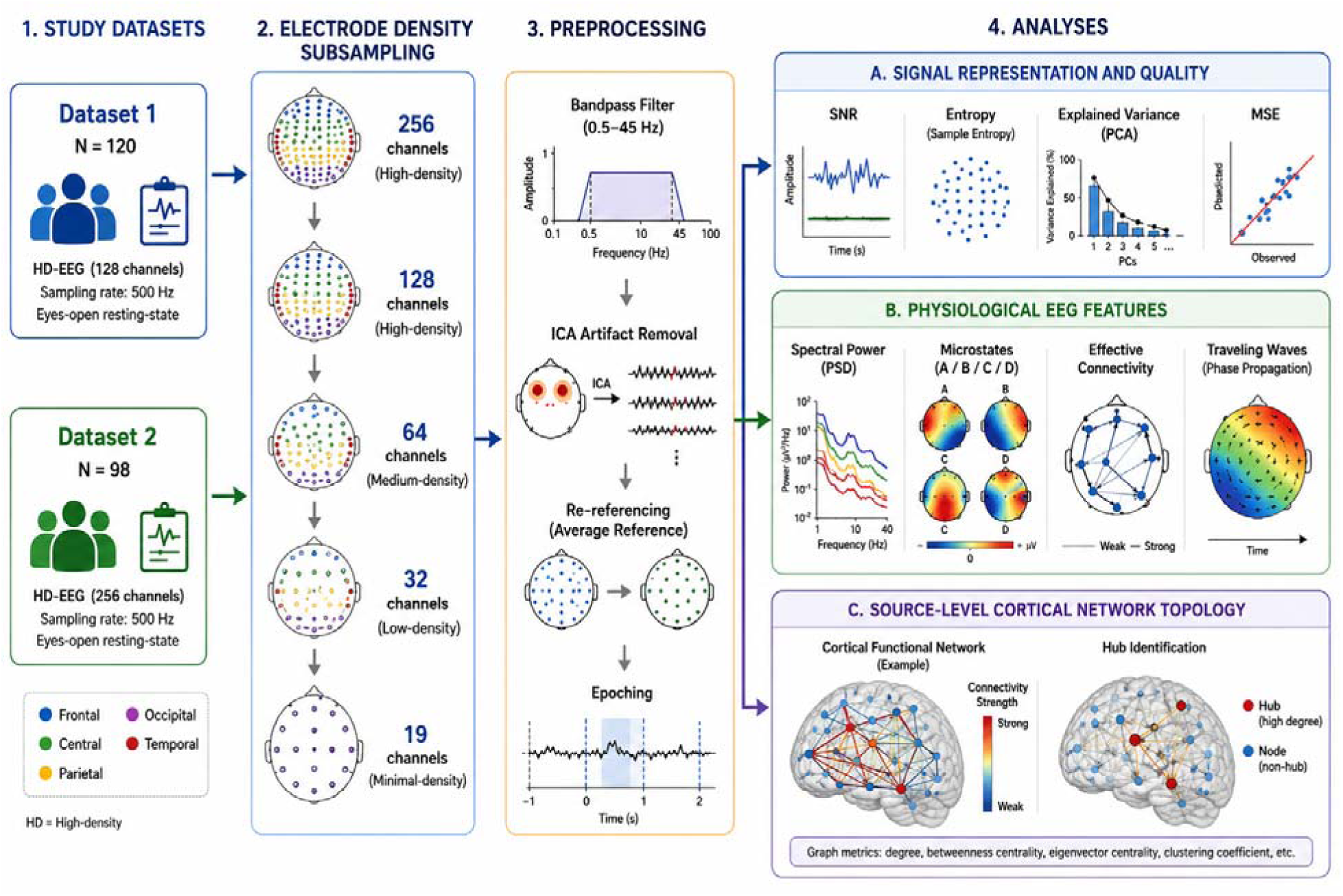
Study design and analytical hierarchy. Two independent high-density resting-state EEG datasets were subsampled to progressively lower electrode-density configurations and processed using an identical preprocessing pipeline. Density effects were then evaluated at three increasingly spatially specific levels: (A) signal representation and quality, including SNR, entropy, explained variance, and MSE (B) physiological scalp-level EEG features, including spectral power, microstates, effective connectivity, and traveling waves; and (C) source-level cortical network topology, assessed through functional connectivity and hub identification.

### 1. Participants and Datasets

Two independent resting-state datasets with different participant characteristics and recording conditions were selected to assess whether density-dependent effects generalize across healthy populations and acquisition settings. Although large-scale open EEG resources have become increasingly available, datasets combining large sample sizes with genuinely high-density recordings remain comparatively scarce, as most large EEG cohorts have been acquired using conventional low or moderate-density systems. This represents an important practical constraint for electrode-density research, because controlled evaluation requires recordings acquired at sufficiently high spatial resolution to allow the same data to be systematically subsampled to multiple lower-density configurations. We therefore selected two comparatively large independent cohorts with original high-density recordings, enabling within-subject comparisons across a broad range of electrode densities while avoiding confounds that would arise from comparing separately acquired low and high-density datasets. The inclusion of two cohorts with distinct population characteristics and acquisition settings further allowed us to examine the reproducibility and generalizability of density-dependent effects across independent empirical data.

**Dataset 1** comprised resting-state EEG recordings from 120 healthy children selected from the Healthy Brain Network (HBN), a large-scale open-science initiative designed to provide multimodal neuroimaging, electrophysiological, behavioral, and phenotypic data across development (Alexander et al., 2017). EEG was acquired at a sampling rate of 500 Hz using a 128-channel HydroCel Geodesic Sensor Net (Electrical Geodesics Inc.) as part of an extensive electrophysiological battery comprising both task-based and task-free paradigms (Langer et al., 2017). The resting-state protocol included alternating eyes-open and eyes-closed periods, and only resting-state data were retained for the present analysis. The resulting sample provided a comparatively large pediatric high-density EEG cohort suitable for systematically evaluating the consequences of spatial subsampling across electrode-density configurations.

**Dataset 2** comprises resting-state EEG recordings from 98 healthy adults originally acquired as part of a study investigating the electrophysiological correlates of real-life creative behavior (Chhade et al., 2024). Participants were recruited from the general population and represented a broad adult age range (18–68 years). Resting-state EEG was recorded for 5–6 min with eyes closed using a 256-channel HydroCel Geodesic Sensor Net at an original sampling rate of 1000 Hz (Chhade et al., 2024). The original study used these recordings to investigate source-level resting-state functional networks associated with individual creativity; in the present work, the resting-state recordings were instead used independently of the behavioral analysis to evaluate the effect of progressive electrode-density reduction. The 256-channel acquisition provides the highest spatial sampling available in the present study and enables controlled subsampling across a broad range of configurations (256, 128, 64, 32, and 19 channels).

#### Ethics and informed consent

The present study involved secondary analyses of two previously collected datasets and did not involve new participant recruitment or data acquisition. Dataset 1 was derived from the Healthy Brain Network (HBN), an open-science resource whose study procedures were approved by the Chesapeake Institutional Review Board. Written informed consent was obtained from the legal guardians of participants younger than 18 years, together with written assent from the participants, and consent procedures included permission for data sharing through IRB-approved protocols (Alexander et al., 2017). Dataset 2 comprised previously acquired data from the CREaNET cohort reported by Chhade et al. (2024). The original study was approved by the ethical committee of the University Hospital Centre of Rennes (agreement No. 20–171), and written informed consent was obtained from all participants. The present work consisted exclusively of secondary analyses of these previously collected data and involved no additional intervention or participant contact.

### 2. Electrode Density Configurations

To simulate progressively reduced electrode densities, the high-density recordings were subsampled into standard configurations spanning the full range from clinical minimal-density to research-grade high-density systems (Figure 2). For Dataset 1 (128-channel maximum), configurations of 128, 64, 32, and 19 channels were used. For Dataset 2 (256-channel maximum), configurations of 256, 128, 64, 32, and 19 channels were used.

**Figure 2.**
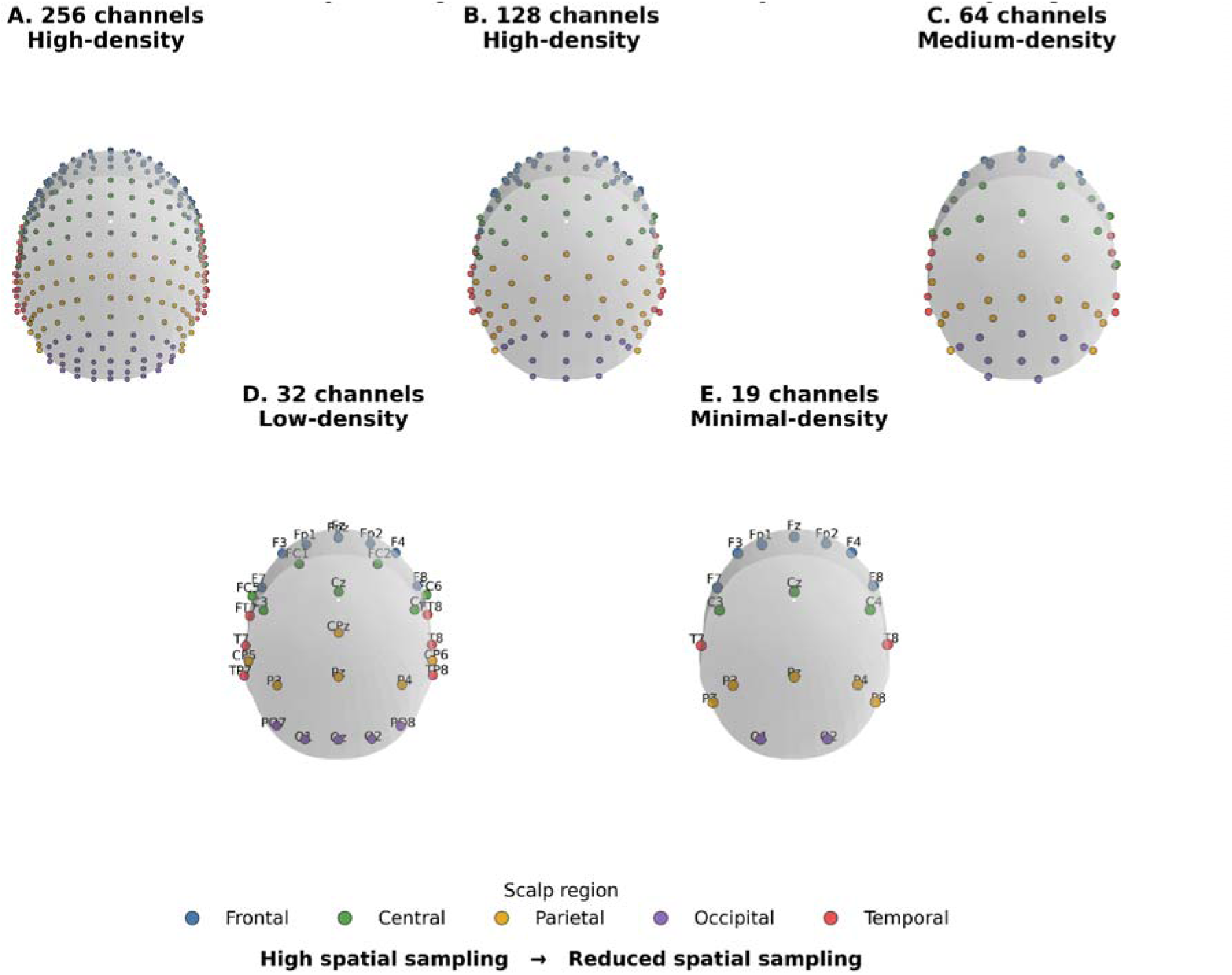
Electrode-density configurations used for spatial subsampling. Standardized scalp representations of the 256, 128, 64, 32, and 19-channel configurations considered across the two datasets. Electrode positions are based on standard 10–5/10–10/10–20 coordinates and are color-coded by the lobe.

Electrode subsets for each configuration were selected by identifying standard 10–5 / 10–10 / 10–20 positions within the full HD cap layout, using nearest-neighbor matching based on 3D electrode coordinates. Please refer toSupplementary materials Table S1 for the electrode names of the different densities.

### 3. Preprocessing

All datasets underwent the same preprocessing sequence independently at each electrode-density configuration to ensure that observed differences reflected spatial sampling rather than a change in processing strategy. The pipeline comprised the following steps:

- Bandpass filtering: 1–45 Hz, using a zero-phase FIR filter. A notch filter was applied at 50 Hz to suppress power line interference.
- Artifact removal: Independent Component Analysis (ICA) was applied to identify and remove components corresponding to ocular (blinks, saccades) and myogenic artifacts (Jung et al., 2000; Delorme & Makeig, 2004). The number of ICA components was set equal to the number of channels minus one for each configuration. Residual artifacts were identified and rejected through visual inspection of the continuous data.
- Re-referencing: All data were re-referenced to the common average across available electrodes within each montage. This step was performed after ICA to ensure the reference reflects only neural signals.
- Epoching: Resting-state data were segmented into non-overlapping 2-second epochs. This epoch length provides sufficient temporal support for estimating conventional resting-state spectral and dynamical measures while allowing multiple approximately stationary segments to be obtained from each recording.
- Subject-level aggregation: All EEG features were first estimated at the individual-subject level. Epoch-level estimates were subsequently averaged within each participant before group-level statistical analyses.

### 4. EEG Analysis Framework

The analytical measures were selected to probe complementary aspects of the EEG signal with progressively greater dependence on spatial sampling. Signal-level measures quantified information quality and reconstruction fidelity independently of a specific neurophysiological interpretation. Physiological scalp-level analyses then assessed whether electrode reduction alters commonly used descriptors of oscillatory activity, large-scale topographic organization, directed interactions, and spatiotemporal propagation. Finally, source-reconstructed connectivity and graph-theoretical analyses were used to determine whether changes in sensor density propagate to anatomically specific network inferences. This multilevel design was intended to distinguish measures that are relatively robust to spatial undersampling from those whose interpretation critically depends on adequate spatial sampling.

#### Level 1 — Signal Representation and Quality

Signal-level metrics were used to determine whether reducing electrode density altered the basic quality, complexity, retained variance, and reconstruction fidelity of the EEG before evaluating physiologically derived features. Four complementary metrics were therefore examined: SNR, entropy, explained variance, and MSE.

##### Signal-to-Noise Ratio (SNR)

SNR was included as a fundamental measure of EEG signal quality because it quantifies the relative contribution of physiologically relevant activity to background noise. SNR was defined as the ratio of signal power in the physiological frequency band (1–45 Hz) to noise power in the high-frequency band (>45 Hz), computed per channel and averaged across all channels within a configuration. Formally:

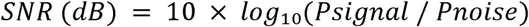

Where *P_signal_* and *P_noise_* are the mean power spectral densities in the respective bands. Higher SNR indicates cleaner signal acquisition with less contamination from non-neural noise sources. SNR was computed per epoch and averaged per subject.

##### Entropy

Entropy was included as a descriptor of signal complexity and irregularity (Lau et al., 2022. Entropy-based measures have been extensively applied to EEG to characterize the regularity, predictability, and complexity of ongoing neural activity across physiological and clinical conditions (Lau et al., 2022; Vecchio et al., 2024). In the present framework, entropy was therefore used to determine whether progressive spatial undersampling alters the complexity of the recorded EEG signal.

Shannon entropy was computed from the amplitude distribution of each channel’s time series:

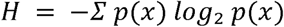

where p(x) is the normalized amplitude histogram. Spectral entropy was additionally computed from the normalized power spectral density across 1–45 Hz. Entropy values were averaged across channels within each configuration. Higher entropy reflects greater signal complexity and a richer representation of ongoing brain activity.

##### Explained Variance

Explained variance was included as a complementary measure of how much of the original EEG variance was retained after ICA-based artifact rejection (Whitmer et al., 2010; Miyakoshi et al., 2014). Here, we used the explained variance ratio to test whether reducing electrode density systematically alters the proportion of EEG variance represented by the retained components after applying the same preprocessing procedure at each density.

Explained variance was computed as the proportion of total signal variance captured by the retained ICA components (i.e., those not classified as artifactual) relative to the total variance of the pre-ICA signal. This metric indexes the effectiveness of artifact removal and the proportion of the signal that is retained for subsequent analysis:

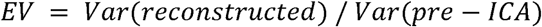

##### Mean Squared Error (MSE)

MSE was selected to provide a direct quantitative measure of the reconstruction error introduced by spatial undersampling. It was used to quantify the reconstruction accuracy of each lower-density configuration relative to the highest available density for that dataset (256-channel for Dataset 2;128-channel for Dataset 1). For each subject and each density, the EEG signal was projected onto the common channel space using spherical spline interpolation, and MSE was computed as:

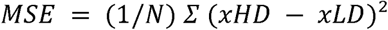

Where xHD and xLD are the HD and lower-density signals at matched electrode positions, and *N* is the number of matched samples. Lower MSE indicates that the reduced configuration more faithfully reproduces the reference HD signal. MSE was included as a direct measure of reconstruction fidelity. By comparing each reduced montage with the corresponding high-density reference after projection to a common electrode space, MSE quantifies the information lost through spatial subsampling rather than characterizing the lower-density signal in isolation.

#### Level 2 — Scalp-level EEG Features

This level examined whether electrode density affects physiologically meaningful descriptors derived directly from multichannel scalp EEG, without introducing the additional modelling assumptions associated with cortical source reconstruction. The selected measures captured complementary dimensions of resting-state activity: spectral power characterized oscillatory content, microstates captured recurrent spatial configurations of the scalp electric field and their temporal organization, MVAR/DTF characterized frequency-specific directed interactions among recorded signals, and traveling-wave analysis quantified the spatiotemporal propagation of oscillatory activity across the scalp.

##### Spectral Power

Spectral power was selected as a benchmark scalp-EEG feature because frequency-domain measures remain among the most widely used quantitative descriptors of resting-state EEG activity, with oscillatory activity conventionally characterized within delta, theta, alpha, and beta frequency bands (Newson & Thiagarajan, 2019; Donoghue et al., 2022). Relative power was examined to quantify the contribution of each band to the overall spectral composition. Quantifying spectral power both globally and across scalp regions further allowed us to test whether electrode reduction preserves overall oscillatory content while altering its spatial distribution.

Relative spectral power was computed per electrode using Welch’s method (Welch, 1967) (window length: 2 s, 50% overlap, Hanning window). Relative power was extracted in four canonical frequency bands: delta (1–4 Hz), theta (4–7 Hz), alpha (8–12 Hz), and beta (13–30 Hz) and was defined as band power normalized by total power across 1–45 Hz. Power values were computed per channel and averaged within standard scalp regions of interest (frontal, central, temporal, parietal, occipital) to assess spatial distribution differences across density configurations.

##### EEG Microstates

EEG microstates have become an established approach for characterizing the temporal organization of spontaneous brain activity during rest (Khanna et al., 2015; Michel & Koenig, 2018). Resting-state EEG microstate analysis was applied to Datasets 1 and 2 to examine whether spatial undersampling alters the characterization of canonical brain microstates.

###### Segmentation

Microstate segmentation was performed independently for each subject and each density configuration using the modified K-means algorithm (Pascual-Marqui et al., 1995). The algorithm clusters EEG scalp topographies at Global Field Power (GFP) peaks (i. e time points at which spatial signal power is locally maximal) using polarity-independent spatial correlation as the assignment criterion. The number of clusters was fixed at k = 4, corresponding to the canonical microstate classes A, B, C, and D established across the literature (Lehmann et al., 1987; Michel & Koenig, 2018). The algorithm was initialized with 20 random restarts, retaining the solution with the highest Global Explained Variance (GEV).

###### Back-fitting

The four cluster templates derived from GFP-peak segmentation were back-fitted to the full continuous time series by assigning each sample to the microstate class whose template map exhibited the highest spatial correlation (polarity-independent) with the instantaneous scalp topography.

Map matching across densities: Because K-means labels are arbitrary, a map-matching step was applied per subject to align microstate labels across density configurations. The four templates from each lower-density solution were matched to the four templates from the highest available density (256-channel for Dataset 2; 128-channel for Dataset 1) using the Hungarian algorithm applied to a pairwise spatial correlation matrix. The quality of each match was quantified using the mean spatial correlation across matched template pairs; subjects with a mean correlation below 0.70 at any density were flagged for review.

###### Temporal parameters

Four parameters were extracted per microstate class per subject per density:

- Duration (ms): mean time a microstate class remains continuously active before switching to another class.
- Occurrence (/s): mean number of times per second the microstate class is activated.
- Coverage (%): proportion of total recording time occupied by each microstate class.
- Transition probabilities (syntax): pairwise transition rates between microstate classes, corrected for occurrence rates to isolate non-random sequential structure (Wackermann et al., 1993).

Additionally, the GEV of the four-class solution was computed per density as a global index of solution quality: the proportion of total EEG variance accounted for by the four microstate templates.

##### Effective Connectivity

Effective connectivity was included to determine whether electrode undersampling affects the inferred direction of interactions among scalp EEG signals. Among directed-connectivity approaches, DTF was selected as it provides a frequency-resolved, multivariate estimate of directed interactions from a single MVAR model fitted jointly to the recorded signals (Kamiński & Blinowska, 1991; Blinowska, 2011). This multivariate formulation is particularly relevant for EEG, where pairwise estimates may identify spurious or indirect interactions when multiple signals are mutually interdependent. Multichannel approaches have been shown to provide more reliable estimates of EEG propagation patterns (Kuś et al., 2004; Blinowska et al., 2004). Directional brain connectivity was assessed using Multivariate Autoregressive (MVAR) modelling followed by computation of the Directed Transfer Function (DTF) (Kaminski & Blinowska, 1991). A model order of 10 was used, selected via the Akaike Information Criterion across subjects. DTF was computed in the frequency bands, providing a measure of directional information flow between scalp regions. Global connectivity was defined as the mean DTF value across all directed electrode pairs within each configuration. Regional connectivity was computed by averaging DTF values within and between predefined scalp lobes (frontal, temporal, parietal, occipital - See table S1 for region affiliation).

##### Traveling Waves

Traveling waves were included as a spatiotemporal descriptor of resting-state EEG. Spontaneous EEG oscillations can organize into propagating phase patterns across the scalp, particularly in the alpha range (Patten et al., 2012; Alamia & VanRullen, 2019; Li et al., 2025). Spatiotemporal propagation of neural oscillations was characterized using phase gradient analysis applied via the Hilbert transform. Instantaneous phase was computed per channel in the theta and alpha bands, and propagation speed was estimated from the spatial gradient of phase differences across the scalp, expressed in meters per second. Delay maps were constructed per lobe by computing the mean phase delay relative to a frontal reference electrode. Traveling wave metrics were computed per epoch and averaged per subject.

#### Level 3 — Cortical Network Topology

This level of analysis tested the most spatially specific inference in the study: whether changing electrode density alters source-reconstructed functional topology and, in particular, the anatomical identity and consistency of cortical hubs. Network hubs at rest have been identified in previous EEG studies using graph-theoretical measures (Jin et al., 2013; Kabbara et al., 2017).

##### Source Reconstruction

Sensor-space EEG signals were projected to cortical source space using the weighted minimum norm estimate (wMNE). Source reconstruction was performed on a standard MNI brain parcellated according to the Desikan-Killiany atlas (68 cortical regions). Source time series were extracted per region of interest.

##### Functional Connectivity and Graph Construction

Pairwise functional connectivity between source-reconstructed regions was computed using the Phase Locking Value (PLV) in the alpha band. Connectivity matrices (68 × 68) were thresholded to retain only significant edges (surrogate-based threshold, p < 0.05), yielding sparse binary graphs for each subject and density configuration.

The wMNE/PLV combination was selected based on previous methodological and comparative studies of EEG source connectivity showing that the choice of inverse solution and connectivity estimator substantially influences the reconstructed network, with wMNE combined with phase-synchronization measures providing good network-reconstruction performance (Hassan et al., 2014, 2017; Allouch et al., 2022, 2023). This combination has also been successfully applied to the reconstruction of resting-state cortical networks and the identification of the hub regions from dense-EEG recordings (Kabbara et al., 2017).

##### Hub Detection

Graph-theoretic measures were computed on the thresholded connectivity matrices using the Brain Connectivity Toolbox (Rubinov & Sporns, 2010).

Functional hubs were defined as nodes ranking in the top 10% of betweenness centrality across subjects. Hub detection was performed separately for each density configuration, and the resulting hub sets were compared across densities.

##### Hub Consistency Analysis

The consistency of hub identification across density configurations was quantified using two complementary metrics:

- Jaccard index: proportion of overlap between hub sets identified at different densities, computed pairwise across all density pairs.
- Spearman correlation: rank-order correlation of betweenness centrality values across all 68 regions, computed between each lower-density configuration and the highest available density.

### 5. Statistical Analysis

All comparisons across density configurations were conducted within subjects (same subjects recorded and reprocessed at all densities), enabling a repeated-measures design that controls for inter-individual variability.

#### Non-Parametric Repeated-Measures Framework

Preliminary normality tests (Shapiro-Wilk) and variance homogeneity tests (Levene) confirmed that the distributions of most metrics violated assumptions of normality and equal variance. Accordingly, all omnibus tests used the Friedman test (non-parametric equivalent of repeated-measures ANOVA) across the five density levels. Significant Friedman results (p < 0.05) were followed up with Wilcoxon signed-rank tests for pairwise density comparisons, with Bonferroni correction applied to control the family-wise error rate.

For signal-level metrics in between-subjects comparisons (where applicable), the Kruskal-Wallis test was used as the omnibus test, followed by Mann-Whitney U tests with Bonferroni correction for pairwise comparisons.

#### Effect Sizes

Effect sizes were reported for all significant comparisons:

- Kendall’s W: for Friedman tests, indicating the proportion of variance explained by the density factor.
- Cohen’s d: for pairwise Wilcoxon comparisons, computed from the standardized mean difference between density conditions.

#### Microstate-Specific Statistics

For microstate temporal parameters, Friedman tests and Wilcoxon signed-rank tests were applied per microstate class (A, B, C, D) and per parameter (duration, occurrence, coverage), with Bonferroni correction accounting for the number of density pairs, microstate classes, and parameters tested simultaneously. For transition probabilities, a chi-square test was used per density to assess deviation from a chance baseline, followed by element-wise Wilcoxon signed-rank tests across densities with Bonferroni correction.

#### Hub Consistency Statistics

Jaccard index values and Spearman correlations were compared across density pairs using Wilcoxon signed-rank tests. A Jaccard index below 0.5 was considered indicative of poor hub overlap, and a Spearman correlation below 0.7 was considered indicative of substantial centrality rank disruption.

### 6. Code and data availability

EEG analyses were performed in MATLAB using established open-source toolboxes together with custom analysis scripts. MVAR modelling, Directed Transfer Function (DTF) estimation, source reconstruction using weighted minimum-norm estimation (wMNE), and source-level Phase Locking Value (PLV) connectivity analyses were implemented using functions from the FieldTrip toolbox (Oostenveld et al., 2011). FieldTrip supports MVAR/source analysis and several connectivity estimators, including DTF and PLV. Absolute and relative spectral power were estimated using MATLAB’s implementation of Welch’s power spectral density method (pwelch; Welch, 1967). Traveling-wave analyses were adapted from the publicly available Travelling-waves-EEG-2.0 toolbox (https://github.com/artipago/Travelling-waves-EEG-2.0), with modifications to accommodate the electrode-density configurations examined in the present study. Microstate analyses were performed using EEGLAB Microstate toolbox, with additional custom scripts for density-wise comparisons of topographies, temporal parameters, and transition matrices. Custom code developed for electrode subsampling, harmonization across density configurations, metric extraction, statistical analyses, and figure generation are made available at https://github.com/AyaKabbara/eeg-density-multilevel.

Dataset 1 is available at http://fcon_1000.projects.nitrc.org/indi/cmi_healthy_brain_network

## Results

### 1. Electrode density has metric-dependent effects on signal representation

#### Signal quality and information content

Across both datasets, SNR increased monotonically with electrode density, whereas entropy showed the same general increase in signal complexity (Figure 3 A). In Dataset 1, density significantly affected SNR (χ²(3) = 24.7, p < 0.001, Kendall’s W = 0.69), with 19- and 32-channel configurations lower than 64 and 128 channels while the 64- and 128-channel configurations did not differ significantly (U = 71.4, p = 0.41, d = 0.29). Dataset 2 replicated this pattern (χ²(4) = 31.2, p < 0.001, Kendall’s W = 0.73): SNR rose from 8.1 dB at 19 channels to 15.8 dB at 256 channels, while 128 and 256 channels did not differ significantly (U = 69.8, p = 0.38, d = 0.21).

**Figure 3.**
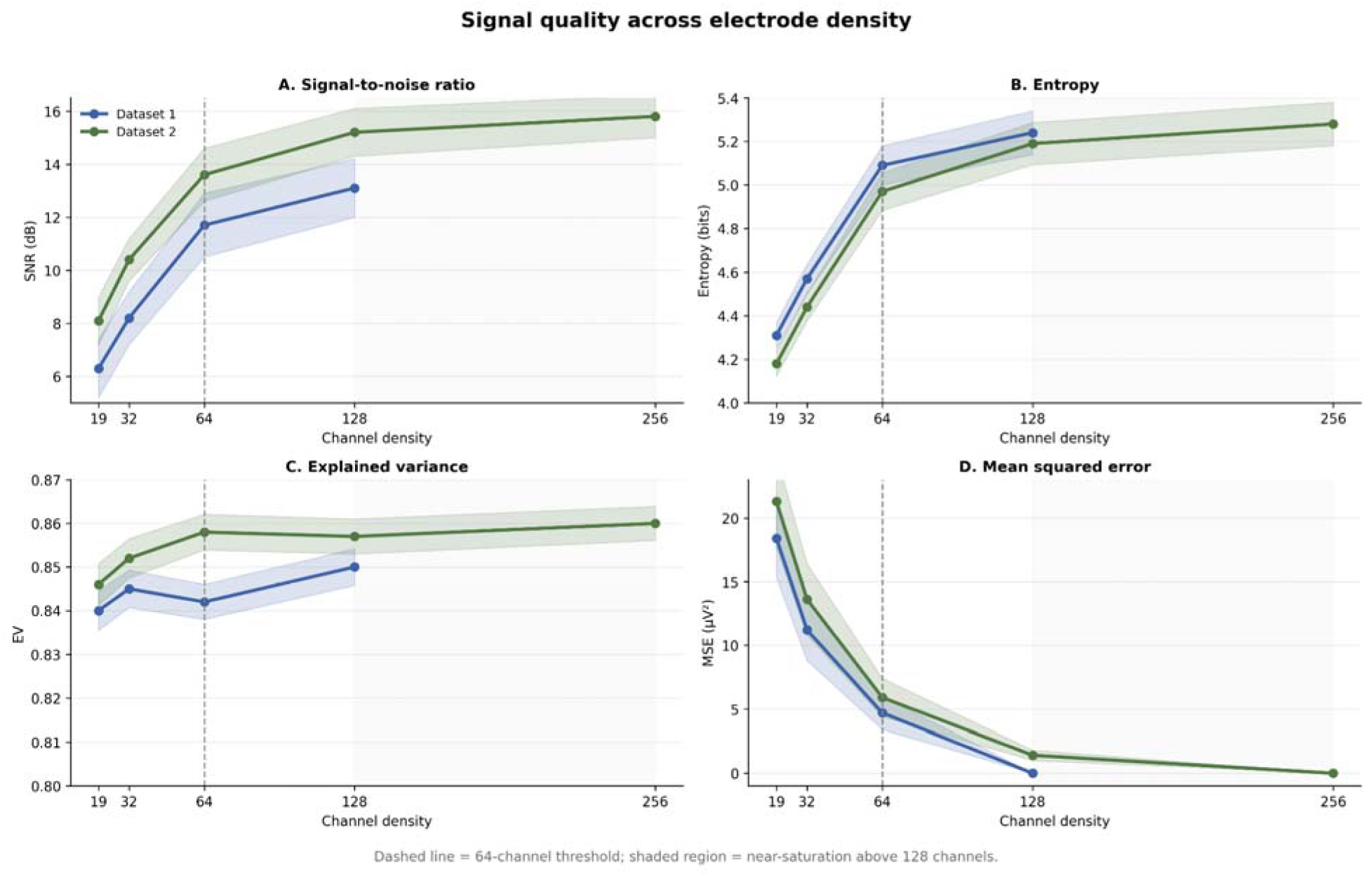
Signal representation across electrode density. SNR, entropy, explained variance, and reconstruction MSE are shown as a function of electrode density for both datasets. The dashed vertical line marks 64 channels as the approximate region in which several signal-level measures began to stabilize.

#### Entropy and retained variance

Entropy results converged across datasets (Figure 3 B). The 19-channel configuration showed lower entropy than higher-density configurations in Dataset 1. 64 and128 channels showed no significant difference in entropy distributions (U = 73.1, p = 0.33, d = 0.24). Dataset 2 showed the same low-density reduction, with no significant difference between 128 and 256 channels (U = 70.9, p = 0.42, d = 0.18). In contrast, explained variance was stable across density in both datasets (Dataset 1: χ²(3) = 2.14, p = 0.52; Dataset 2: χ²(4) = 3.61, p = 0.47), with median values between 0.81 and 0.86 (Figure 3 C). Thus, reducing density did not uniformly degrade every signal descriptor.

#### Reconstruction error

MSE provided the complementary reconstruction perspective: error decreased monotonically as density increased (Figure 3 D). In Dataset 1, 19- and 32-channel configurations differed significantly from 64 channels, whereas 64 and 128 channels did not (U = 68.2, p = 0.31, d = 0.27). In Dataset 2, the same low-density penalty was observed, while 128 and 256 channels were similar (U = 72.6, p = 0.29, d = 0.22). Together, the signal-level results indicate that spatial undersampling changes signal complexity and reconstruction fidelity more strongly than it changes the proportion of variance retained after preprocessing.

### 2. Global spectral information is preserved, whereas regional EEG features are density-sensitive

#### Spectral Power

Global relative spectral power was preserved across electrode densities in both datasets (all Friedman p > 0.05 across frequency bands), indicating that broad oscillatory composition can be recovered even from sparse montages. Regional power estimates were more density-sensitive. Frontal relative power differed across densities in both datasets (Dataset 1: χ²(3) = 17.3, p < 0.01 -see Figure 4 A; Dataset 2: χ²(4) = 21.8, p < 0.001 - see Figure 4 B), and temporal and parieto-central distributions also shifted as channel number decreased. Thus, the overall spectrum remained stable while its apparent spatial distribution changed.

**Figure 4.**
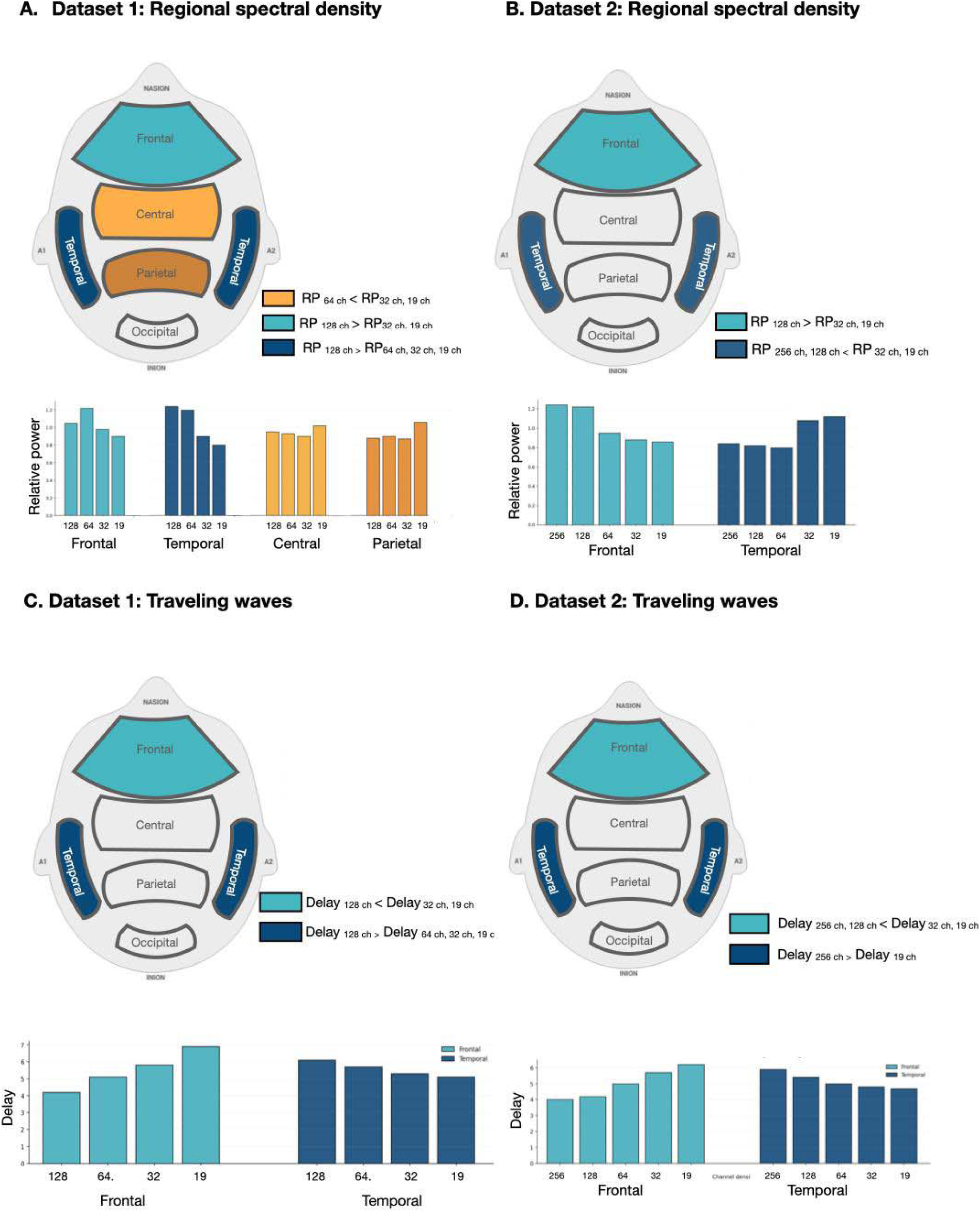
Density effects on regional spectral power and traveling-wave estimates. Regional relative spectral power and phase-propagation measures are shown for Datasets 1 and 2 across electrode-density configurations. Bar plots summarize the corresponding regional values across channel counts.

#### Traveling Waves

Traveling-wave estimates showed stronger density dependence (Figure 4 C, Figure 4 D). Propagation speed was significantly lower at 19 and 32 channels than at 64 channels (χ²(3) = 16.78, p < 0.01, W = 0.47; 19 vs. 64: U = 49.7, p < 0.0025, d = 1.08; 32 vs. 64: U = 53.2, p < 0.0025, d = 0.74). Regional delay estimates showed the same sensitivity: frontal and temporal delays showed significant differences between high and low densities. No significant difference in wave speed or delay was observed between 128 and 256 channels. For detailed statistical results, please refer to the supplementary materials.

#### Effective Connectivity

Effective connectivity showed a clear density gradient in both datasets (Figure S3). Global DTF decreased as electrode density was reduced, with the strongest decline at lower channel counts. In Dataset 1, 128 channels exceeded 64, 32, and 19 channels, and 64 channels exceeded both 32 and 19 channels (all reported pairwise p < 0.0025). Dataset 2 showed the same overall pattern: 128 and 256 channels exceeded 64 channels and below, whereas 128 and 256 channels did not differ significantly (U = 71.3, p = 0.33, d = 0.23).

#### Regional connectivity

Regional connectivity estimates were also density-dependent (Figure 5). Dataset 2 showed stronger intra-frontal and intra-temporal connectivity at 128–256 channels than at lower densities (χ²(4) = 21.8, p < 0.001), while Dataset 1 showed changes in intra-temporal and temporo-parietal connectivity across densities (χ²(3) = 17.3, p < 0.01). The direction of individual regional effects was not identical across cohorts, but both datasets showed that changing electrode density alters the apparent spatial organization of directed connectivity.

**Figure 5.**
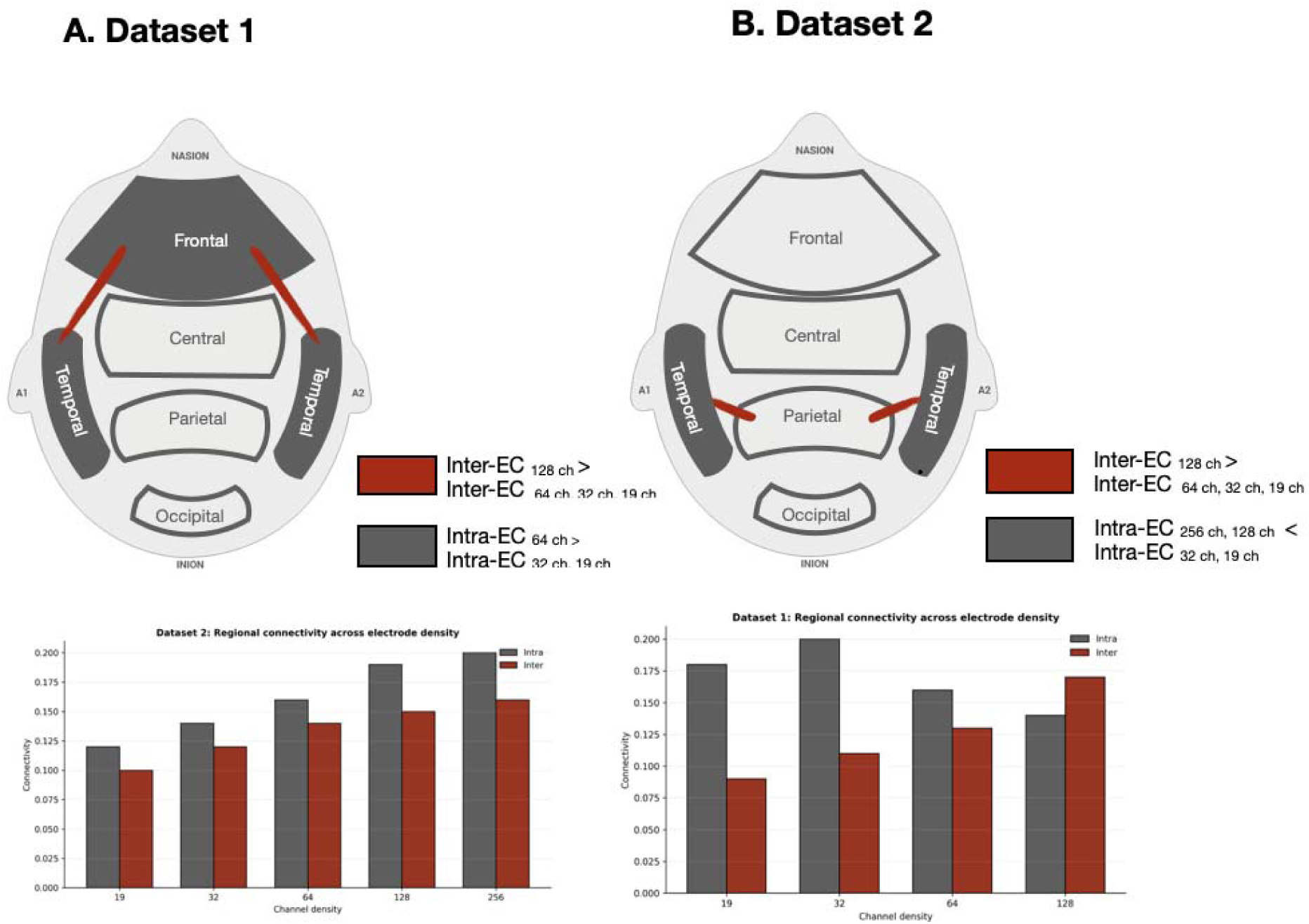
Regional effective connectivity across electrode density. Schematic scalp-region summaries and accompanying bar plots show density-dependent changes in intra-regional and inter-regional directed connectivity for Datasets 1 and 2.

### 3. Canonical microstate classes persist, but their temporal characterization is density-sensitive

All four canonical microstate classes (A–D) were returned at every density because the clustering solution was fixed at k = 4. The relevant question was whether their topographies and derived temporal parameters remained consistent with the high-density reference. Figure 6 shows the results obtained in microstate analysis computed on Dataset 1 (see Figure S2 for dataset2 results). Map-matching correlations exceeded 0.70 for most subjects at all densities (93.1% in Dataset 1; 95.4% in Dataset 2), indicating broad preservation of class identity despite progressive changes in quantitative microstate characteristics.

**Figure 6.**
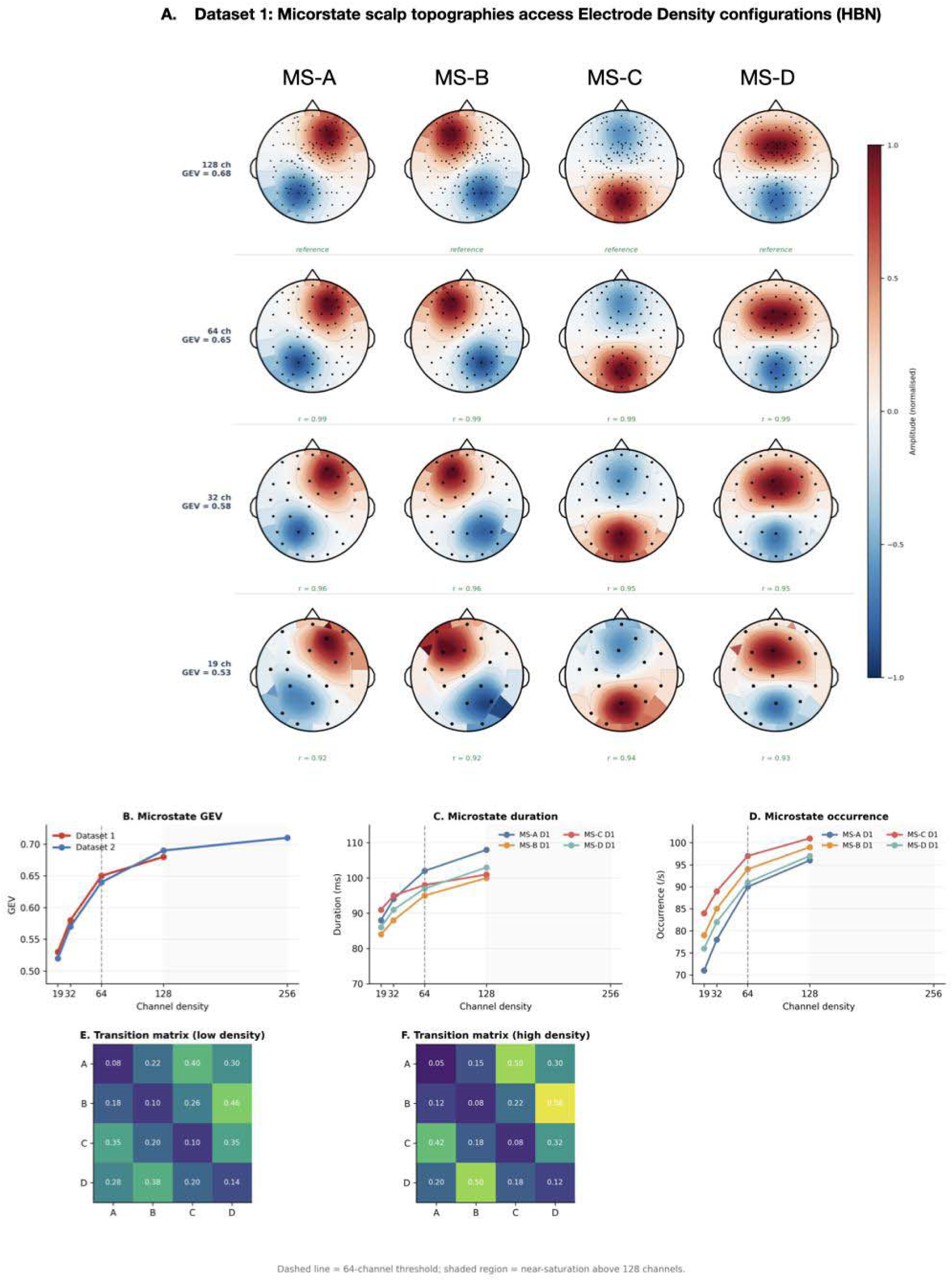
Microstate topography and dynamics across electrode density. For Dataset 1, canonical microstate maps A–D are shown across 128-, 64-, 32-, and 19-channel configurations, together with density-dependent changes in global explained variance (GEV), duration, occurrence, and transition structure. The corresponding Dataset 2 results are provided in the Supplementary Material.

#### Microstate fit and temporal parameters

GEV decreased as electrode density was reduced in both datasets. In Dataset 1, median GEV declined from 0.68 at 128 channels to 0.53 at 19 channels (χ²(3) = 19.8, p < 0.001, W = 0.55), while Dataset 2 showed a similar decline from 0.71 at 256 channels to 0.52 at 19 channels (χ²(4) = 25.6, p < 0.001, W = 0.60). In both cohorts, the highest densities were similar, whereas 19- and 32-channel solutions showed the largest reductions (Figure 6 B).

#### Duration and occurrence

Microstate duration shortened with decreasing density, while occurrence increased. At high density, mean durations across classes were approximately 80–110 ms; at 19 channels they were 18–26 ms shorter. Dataset 1 showed significant low-versus high-density differences for several classes, particularly MS-A and MS-D (Figure 6 C), and Dataset 2 replicated the overall duration gradient with no significant 128-versus-256 differences (Figure). Occurrence showed the complementary pattern, consistent with greater fragmentation of microstate segments at lower density (Figure 6 D).

#### Coverage and transition structure

Coverage was comparatively stable, but class-specific changes were observed. MS-A coverage decreased by an estimated 6.8 percentage points from high density to 19 channels, while MS-D increased by approximately 5.9 percentage points. Dataset 1 showed lower MS-A coverage at 19 and 32 channels than at 64 and 128 channels, whereas Dataset 2 did not show significant coverage differences across densities.

Non-random microstate syntax remained detectable at every density (p < 0.001 against the occurrence-corrected baseline in both datasets), but the detailed transition structure became less similar to the high-density solution as density decreased.

At 128–256 channels, the canonical A↔C and B↔D transition asymmetries were clearly expressed. These asymmetries weakened progressively at lower densities, with significant differences for A→C, C→A, B→D, and D→B in comparisons involving 19 channels (all reported p < 0.0025). The 64-channel configuration remained within a non-significant margin of the high-density transition structure in both datasets. Thus, low-density EEG preserved the broad existence of recurrent microstate classes more effectively than it preserved their quantitative temporal organization.

### 4. Spatial undersampling progressively alters cortical network inference

#### Hub identity and consistency

Hub identity became progressively less stable as density decreased. In Dataset 1, the 128-channel solution included canonical resting-state hubs such as posterior/isthmus cingulate, rostral anterior cingulate, parahippocampal, and middle orbitofrontal regions. At 64 channels some canonical hubs were less consistently recovered, while 32 and 19 channels increasingly introduced atypical regions (Figure S3). Dataset 2 showed the same qualitative progression across the broader 256-to-19-channel range: 256 and 128 channels produced the most mutually consistent hub sets, whereas the lowest densities showed the greatest divergence (Figure 7 A).

**Figure 7.**
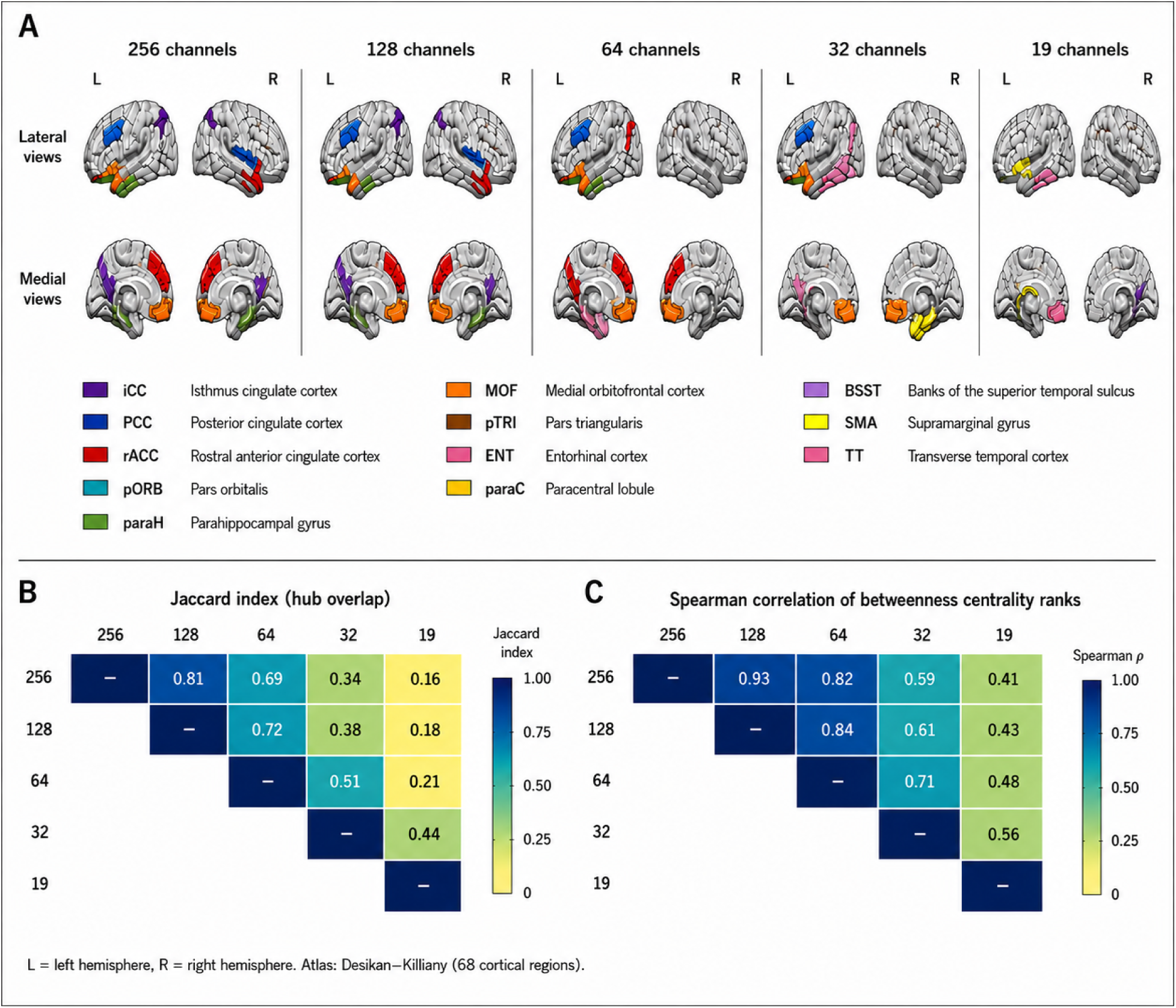
Effect of electrode density on cortical hub identification and cross-density consistency. (A) Cortical regions identified as functional hubs across the 256-, 128-, 64-, 32-, and 19-channel configurations. Hubs were defined as regions belonging to the top 10% of betweenness-centrality values in source-level functional connectivity networks reconstructed on the 68-region Desikan–Killiany cortical atlas. Lateral and medial views of both hemispheres are shown, with colors indicating the anatomical identity of detected hubs. (B) Pairwise consistency of hub identification across electrode-density configurations. The Jaccard index quantifies overlap between the sets of detected hubs, whereas Spearman’s ρ quantifies preservation of the rank ordering of regional betweenness centrality across all cortical regions. L, left hemisphere; R, right hemisphere.

Quantitative consistency analyses confirmed this pattern. Pairwise Jaccard overlap and Spearman correlation of centrality ranks were highest between the densest configurations and declined as lower-density montages were compared. Across the combined comparisons, 256 versus 128 channels yielded Jaccard = 0.81 and ρ = 0.93, while 128 versus 64 remained acceptable (Jaccard = 0.72; ρ = 0.84). Agreement deteriorated markedly for 32- and 19-channel configurations. For example, 256 versus 19 channels yielded Jaccard = 0.16 and ρ = 0.41. Comparisons involving densities below 64 channels showed the weakest agreement.

These results indicate that the strongest effect of spatial undersampling occurs when the scientific claim depends on the anatomical identity or rank ordering of source-level network nodes, rather than on broad global signal properties.

### 5. Convergence across analyses reveals a hierarchy of density sensitivity

Across the two independent datasets, electrode-density effects followed a coherent hierarchy rather than a uniform loss of EEG information. Broad global descriptors such as spectral power and explained variance were comparatively preserved. Regional spectral estimates, microstate dynamics, traveling waves, and effective connectivity became progressively more sensitive to density. Source-level hub topology showed the strongest loss of agreement at sparse montages. Although the exact transition depended on the metric, many spatially dependent measures showed marked degradation below approximately 64 channels, whereas 128- and 256-channel estimates were generally similar.

## Discussion

This study shows that electrode density does not exert a uniform effect across the EEG analysis chain. Measures that summarize activity over the whole montage, particularly global relative spectral power and retained variance after preprocessing, were comparatively stable. Sensitivity increased when the analysis required a spatial distribution, a phase gradient, a directed interaction, or the anatomical identity of a source-level node. This hierarchy was reproduced in two independent cohorts despite differences in age, acquisition system, and maximum channel count. The principal implication is therefore not that one montage is optimal for every EEG application, but that the minimum adequate density depends on the spatial specificity of the intended inference. Results are discussed hereafter.

### Interpreting the approximately 64-channel transition

Across the present analyses, 64 channels repeatedly marked the range above which several estimates began to stabilize and below which deviations from the high-density reference became more pronounced. This should be described as an empirical transition observed under the current resting-state conditions, rather than as a biological cutoff. It nevertheless converges with several independent lines of evidence. Hatlestad-Hall et al. (2023) subsampled 128-channel resting EEG to 64, 32, and 19 electrodes and evaluated four inverse solutions, four connectivity measures, and five graph metrics. Agreement with the 128-channel reference decreased as density fell, and the authors recommended at least 64 electrodes for source-level graph analysis. In simulation, Allouch et al. (2023) similarly found that at least 64 channels were required for accurate reconstruction of cortical resting-state networks, although performance also depended on the inverse solution and connectivity estimator. Liu et al. (2018), using 256-channel recordings reduced to 128, 64, and 32 channels, showed that montage density strongly affected the recovery of source-space resting-state networks, particularly distributed networks containing deeper regions. These findings provide a direct context for the sharp decline in hub overlap and centrality-rank agreement observed here below 64 channels.

The 64-channel result is also compatible with the broader source-localization literature, in which denser and more complete scalp coverage generally reduces localization error and spatial aliasing (Srinivasan et al., 1998; Grieve et al., 2004; Song et al., 2015). However, these studies do not support a universal numerical rule. The spatial Nyquist analysis of Srinivasan et al. (1998), for example, argued that approximately 128 sensors may be needed to sample the full adult scalp field adequately, whereas empirical source-network studies often identify 64 channels as a good compromise between fidelity and feasibility. The apparent discrepancy reflects different endpoints: complete sampling of the scalp potential field is a stricter objective than achieving acceptable stability for a particular derived measure. In the present data, the similarity of many 128- and 256-channel estimates suggests diminishing returns for the outcomes tested, while the remaining advantage of the denser arrays for hub identity cautions against treating 64 channels as equivalent to high-density EEG.

### Global preservation does not imply regional equivalence

The contrast between global and regional results is central to the interpretation of low-density EEG. Averaging spectral power over the montage can preserve the overall oscillatory composition even when the sampled topography is incomplete. Regional estimates, by contrast, depend on which electrodes represent each region and on how well they sample local spatial gradients. This explains why global relative power was stable while frontal, temporal, and parieto-central distributions changed with density. Earlier work likewise showed that electrode density can influence scalp power maps and that dense recordings introduce practical problems such as electrolyte bridging that must be controlled rather than assumed away (Greischar et al., 2004). Thus, additional electrodes are beneficial only when they provide independent, well-distributed measurements of adequate quality.

Frontal and temporal findings deserve particular caution. These regions were repeatedly density-sensitive, but the direction of the temporal effect was not consistent across cohorts: temporal values increased at low density in Dataset 1 and decreased in Dataset 2. The robust finding is therefore a loss of regional stability, not a systematic increase or decrease attributable to low density. Several mechanisms may contribute. Sparse montages change the number, location, and scalp coverage of sensors entering each regional average. The reduction is especially consequential near the inferior and lateral head surface, where temporal coverage is often limited. Volume conduction, edge effects, muscle and ocular contamination, and changes in the common-average reference can then redistribute activity differently across montages. Cohort differences in age, head geometry, resting protocol, and hardware may further determine the sign of the bias. Individual-level analyses may reveal additional heterogeneous patterns that cancel or reverse at the group level. Accordingly, frontal or temporal conclusions derived from 19- or 32-channel resting EEG should be supported by sensitivity analyses across electrode subsets, subject-level estimates, and preferably an independent montage or dataset.

### Signal Quality and Spatial Information

At the signal level, the results suggest an important distinction between signal cleanliness and spatial information content. Measures such as SNR, entropy, and MSE degraded significantly below 64 channels, while explained variance remained relatively stable across densities. This pattern suggests that lower-density systems are not necessarily recording more noise in a simple sense; rather, they are recording a spatially impoverished version of the neural signal. The recorded activity remains broadly interpretable, but it loses the spatial resolution needed to attribute activity to specific scalp regions or underlying cortical generators with confidence.

This means that low-density EEG may still be acceptable for some global applications, such as coarse spectral characterization, while being less suitable for analyses that rely on fine spatial distinctions. Put differently, the signal is still present, but the map of that signal is less detailed. That is why global power may remain relatively preserved even when regional distributions become less stable.

### Microstate topology is more robust than microstate quantification

The microstate results refine, rather than contradict, previous density studies. Zhang et al. (2021) compared 91-, 64-, 32-, 19-, and 8-channel configurations during baseline and propofol sedation. Microstate parameters were broadly consistent across 91, 64, and 32 channels, whereas most discrepancies appeared after including 19 or 8 channels. Condition differences detected at 91 channels were retained at 64 and 32 channels but disappeared at the sparsest densities. In the present study, the broad identity of the four matched classes was likewise preserved across montages, yet GEV, duration, occurrence, coverage for selected classes, and transition structure were not equally stable. This convergence suggests that a sparse montage may recover recognizable canonical maps while still biasing the numerical features subsequently used as biomarkers.

Our findings also extend prior work in three ways. First, they were reproduced in two relatively large independent resting-state cohorts rather than within a single pharmacological transition. Second, they show that different microstate outputs have different density requirements: class correspondence was more robust than temporal segmentation and syntax. Third, they place microstate sensitivity within the same pipeline as signal, spectral, propagation, connectivity, and source-network measures. Shorter durations and more frequent occurrences at low density are consistent with fragmentation of the fitted sequence, but this mechanism should not be viewed as the only explanation. Reduced topographic resolution can make adjacent samples switch more readily between similar templates, while density-specific referencing and clustering can also modify the maps used for back-fitting. The fixed four-class solution further guarantees that four classes are returned; it does not demonstrate that four is the optimal model order at every density. Future work should therefore compare data-driven model order, fixed versus common group templates, and cross-density back-fitting to determine which step produces the observed temporal bias.

### Frequency-specific scope

The emphasis on theta and alpha for traveling waves and on alpha for source connectivity was motivated by the prominence of these rhythms during spontaneous activity and by their established role in large-scale resting-state organization. Alpha is typically the dominant eyes-closed rhythm and has been repeatedly linked to resting network topology, while theta can support slower large-scale propagation. Prior work from our group also found meaningful and broadly interpretable connectivity patterns across multiple canonical bands in resting EEG, (Kabbara et al., 2017). This supports the usefulness of band-limited analysis, but it does not establish that density effects are identical across frequency.

Spatial sampling requirements may vary with the spatial scale, signal-to-noise ratio, and generators of each rhythm. Allouch et al. (2023) demonstrated that density effects on reconstructed networks interact with the connectivity method, and clinical work has reported that density-dependent cognitive associations may be most visible in alpha and beta bands (Dattola et al., 2026). Higher-frequency beta or gamma activity may be more spatially focal and more vulnerable to muscle contamination, whereas slower delta and theta activity may be more spatially diffuse. These expectations require direct testing rather than extrapolation. A complete frequency-sensitivity analysis should therefore evaluate delta, theta, alpha sub-bands, beta, and—where acquisition quality permits—gamma, using the same cross-density framework and correction strategy.

### Hub Detection and Network Topology

Hub detection is perhaps the clearest example of how low-density EEG can remain formally interpretable while becoming increasingly uncertain in anatomical terms. At higher densities, the detected hubs were more consistent with canonical default mode network regions, including posterior cingulate, isthmus cingulate, rostral anterior cingulate, and parahippocampal areas, which aligns with resting-state network literature and source-reconstructed EEG findings (Kabbara et al., 2018,2021; Buckner et al., 2008; Stoyell et al., 2021). As density decreased, however, the set of detected hubs became less stable and more likely to include atypical regions.

The Jaccard and Spearman consistency analyses support this interpretation. Hub overlap and rank order were relatively well preserved at higher densities, but agreement declined sharply below 64 channels. That means the network topology is not simply “present” or “absent” in low-density EEG; instead, the reliability of the inferred topology changes gradually as spatial sampling worsens. For this reason, hub maps derived from 32- or 19-channel systems should be treated cautiously, especially when the specific anatomical identity of hubs is part of the scientific claim (Stoyell et al., 2021; Yamazaki et al., 2013).

### A multilevel framework, not an exhaustive measurement set

The selected outcomes were intended to sample several levels of inference rather than to exhaust the EEG feature space. Signal representation was indexed by SNR, entropy, retained variance, and reconstruction error. In addition, spontaneous physiology was quantified by spectral power, microstates, directed connectivity, and traveling waves; and anatomical network inference by source connectivity and hubs. Other measures that were not examined here may respond differently. At the signal level, one could examine channel-wise correlation, covariance structure, aperiodic spectral parameters, peak alpha frequency, or artifact-detection performance. At the feature level, phase-amplitude coupling, coherence, imaginary coherence, phase-lag indices, mutual information, complexity measures, and microstate map dissimilarity may reveal distinct sensitivities. At the network level, density may affect strength, clustering, efficiency, modularity, participation coefficient, rich-club organization, and dynamic connectivity in different directions. Indeed, Hatlestad-Hall et al. (2023) found that reduced density overestimated mean strength and clustering while underestimating path length, illustrating why one graph metric cannot stand for network validity as a whole. In this study, the convergence across several complementary outcomes is therefore informative, but it should be interpreted as a framework for validation rather than a comprehensive certification of any montage.

### Resting-state boundaries and implications for task and clinical EEG

Both datasets contained healthy participants at rest. Resting-state activity is continuous, spontaneous, and often dominated by spatially broad oscillations; task-evoked and clinical signals can impose different requirements. Event-related potentials may remain detectable at a small set of electrodes when the component is large and the region and time window are specified a priori, yet sparse sampling may alter peak topography, source localization, multivariate decoding, or the statistical specificity of regional effects. Recent ultra-high-density work on visual processing showed that denser local sampling improved category decoding, topographic detail, and localization of the N170 generator (Schreiner et al., 2026). Clinical abnormalities may be either diffuse, making global low-density markers feasible, or focal and deep, making coverage and source modeling critical. In an exploratory MCI cohort, Dattola et al. (2026) found clearer cognition-network associations, stronger group discrimination, and better classification with 173 than with 64 or 18 channels. These observations argue against transferring the present resting-state threshold directly to ERP, BCI, epilepsy, or patient-monitoring applications.

An important next step is to test whether electrode reduction preserves effects rather than only values. This includes group contrasts, within-subject task modulations, correlations with behavioral performance or symptom severity, diagnostic classification, and longitudinal change. A montage may yield an acceptable global mean but attenuate a brain-behavior correlation or move a regional maximum enough to change the scientific conclusion. Density validation should therefore be tied to the inferential target of the study.

### Methodological Considerations

#### Several design choices constrain interpretation

First, lower-density datasets were created by subsampling the same high-density recordings. This is a strong control for neural state and inter-individual variability, but it does not reproduce all characteristics of a physically separate low-density system, including cap fit, sensor technology, impedance distribution, amplifier properties, preparation time, and participant movement.

Second, preprocessing was repeated at each density. The number of ICA components, channels contributing to the average reference, rejected or interpolated sensors, and residual artifact structure therefore changed with montage size. These are realistic consequences of using different systems, but they mean that the observed effect is the combined influence of spatial sampling and its downstream preprocessing consequences.

Third, the two cohorts improve external validity but are not a formal replication under identical conditions. They differed in age, equipment, original density, and resting protocol. These differences may explain why some regional effects agreed in location but not direction. Fourth, regional sensor averages were based on unequal and density-dependent electrode sets. Results may therefore reflect both reduced channel count and altered spatial coverage. Future studies should compare alternative subset geometries at the same channel count and quantify regional coverage or inter-electrode distance directly.

Fifth, source results were obtained with one inverse method, one connectivity measure, one atlas, and a specific graph-thresholding and hub definition. Because inverse, connectivity, leakage correction, parcellation, and threshold choices interact with density (Mahjoory et al., 2017; Allouch et al., 2023; Hatlestad-Hall et al., 2023), robustness should be evaluated across an analytic multiverse rather than assumed from a single pipeline.

Finally, group-level averages do not establish that the same density is adequate for every participant. Head size and shape, skull conductivity, electrode positioning, data quality, dominant rhythm, and source configuration can all modify spatial sampling requirements. Test-retest data and individual error curves would allow a probabilistic recommendation. Such analyses would be especially valuable before low-density measures are used for individual diagnosis or treatment monitoring.

### Practical implications and conclusions

Electrode density should be selected from the endpoint backward. Nineteen- or 32-channel systems may be adequate for broad global spectral summaries when portability, cost, preparation time, or participant tolerance are decisive. They are less secure for regional frontal or temporal claims, microstate temporal biomarkers, traveling-wave gradients, directed regional connectivity, and the anatomical identity of cortical hubs. In the present resting-state datasets, 64 channels represented a pragmatic minimum for many spatially dependent measures, while 128 channels generally provided greater stability and 256 channels produced limited additional gains for several outcomes. This ordering is guidance for comparable recordings and pipelines, not a universal rule.

Overall, spatial undersampling does not simply remove EEG information. It changes which conclusions remain defensible. Global summaries can appear stable while regional distributions, temporal state parameters, and source-network topology are altered. The inconsistent direction of some regional effects across cohorts is itself informative, suggesting that sparse sampling can introduce context- and participant-dependent bias rather than a predictable offset. Claims based on low-density frontal and temporal activity or source-level topology should therefore be accompanied by montage-specific validation and sensitivity analyses. Matching electrode density to the spatial scale of the hypothesis is more scientifically meaningful than classifying EEG systems as globally adequate or inadequate.

**Table 1.**
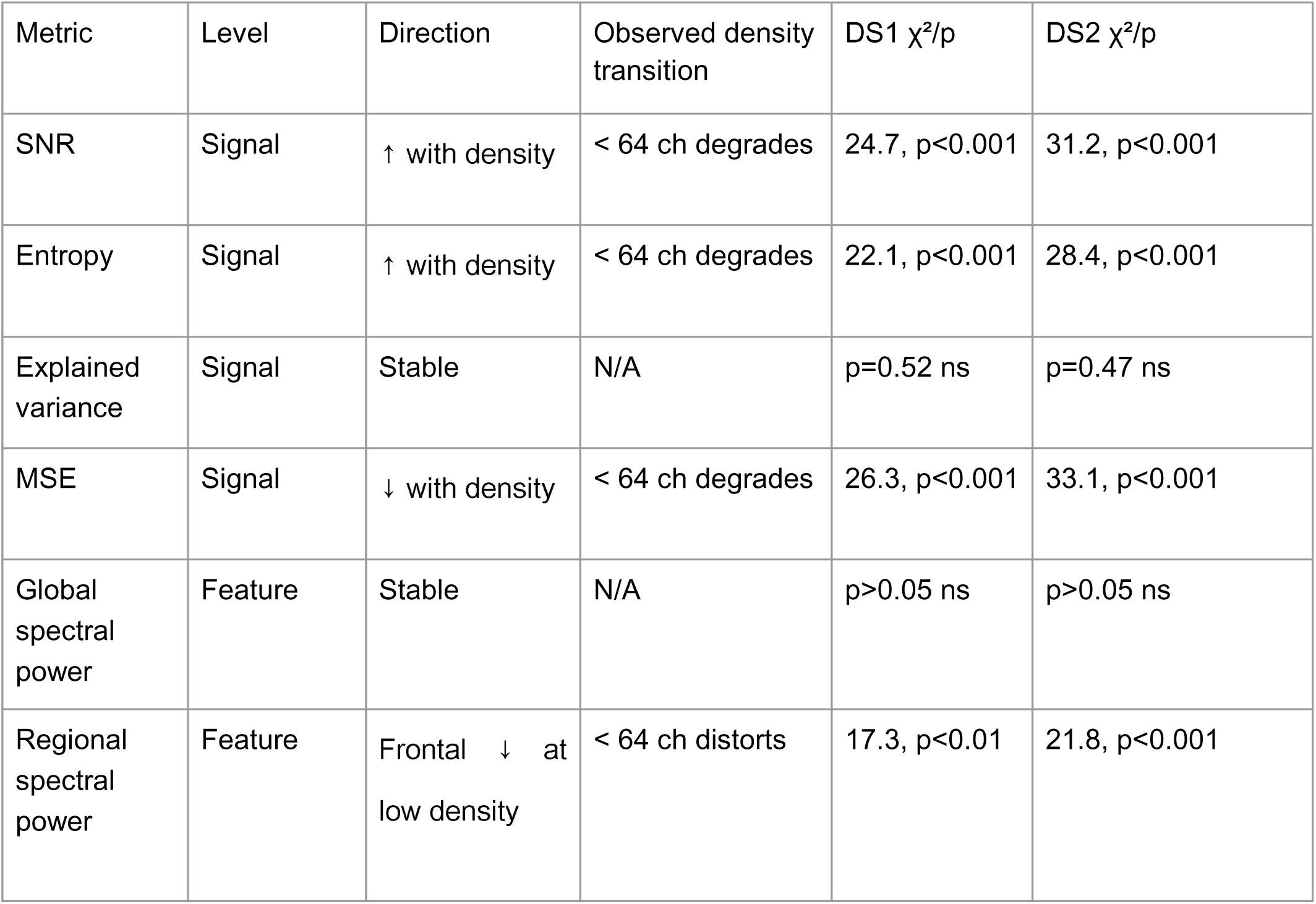
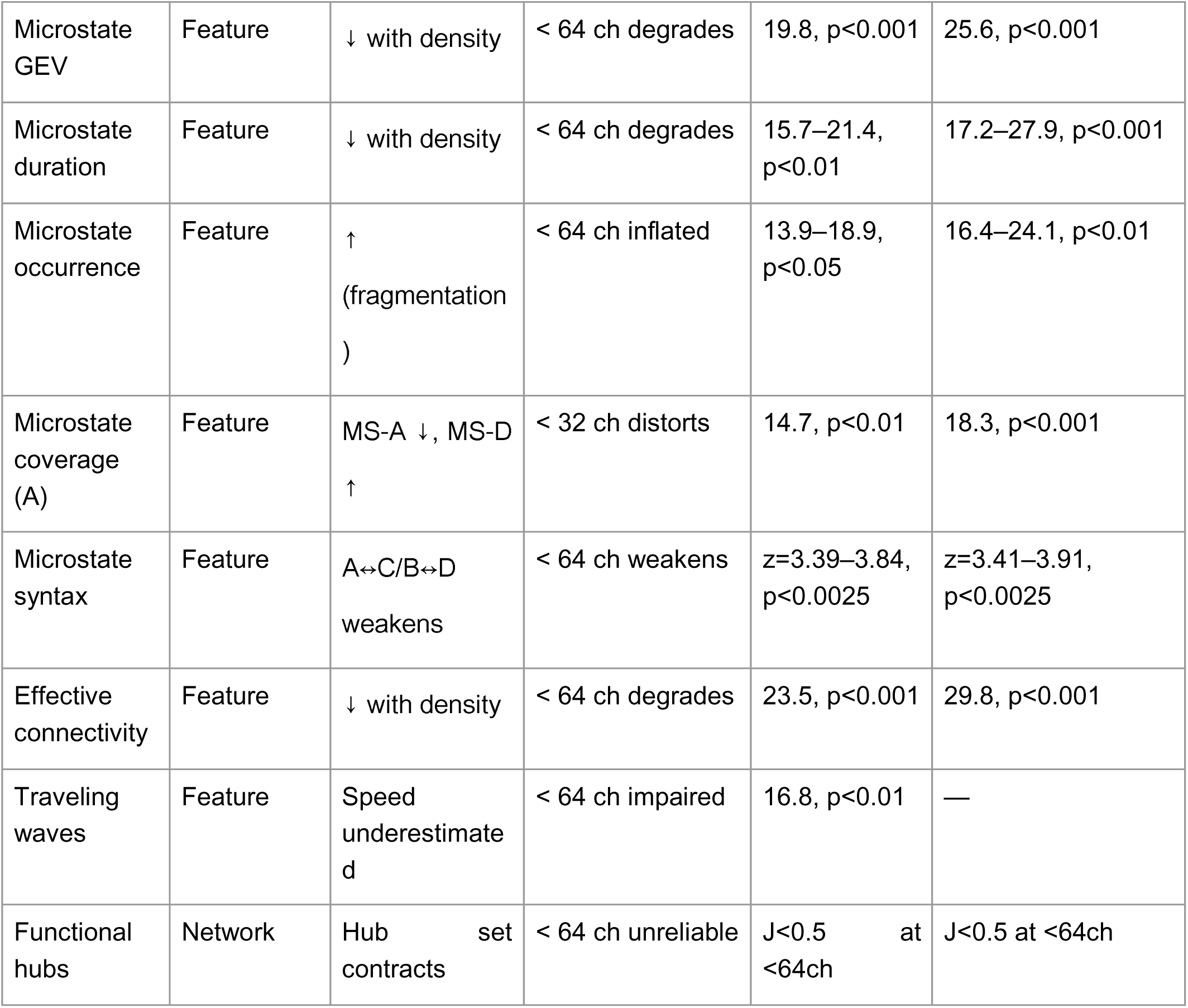
Summary of density effects across signal-level, feature-level, and cortical network measures.

| Metric | Level | Direction | Observed density transition | DS1 $\chi^2/p$ | DS2 $\chi^2/p$ |
| --- | --- | --- | --- | --- | --- |
| SNR | Signal | ↑ with density | < 64 ch degrades | 24.7, $p<0.001$ | 31.2, $p<0.001$ |
| Entropy | Signal | ↑ with density | < 64 ch degrades | 22.1, $p<0.001$ | 28.4, $p<0.001$ |
| Explained variance | Signal | Stable | N/A | $p=0.52$ ns | $p=0.47$ ns |
| MSE | Signal | ↓ with density | < 64 ch degrades | 26.3, $p<0.001$ | 33.1, $p<0.001$ |
| Global spectral power | Feature | Stable | N/A | $p>0.05$ ns | $p>0.05$ ns |
| Regional spectral power | Feature | Frontal ↓ at low density | < 64 ch distorts | 17.3, $p<0.01$ | 21.8, $p<0.001$ |
| Microstate GEV | Feature | ↓ with density | < 64 ch degrades | 19.8, p<0.001 | 25.6, p<0.001 |
| Microstate duration | Feature | ↓ with density | < 64 ch degrades | 15.7–21.4, p<0.01 | 17.2–27.9, p<0.001 |
| Microstate occurrence | Feature | ↑<br>(fragmentation) | < 64 ch inflated | 13.9–18.9, p<0.05 | 16.4–24.1, p<0.01 |
| Microstate coverage (A) | Feature | MS-A ↓, MS-D ↑ | < 32 ch distorts | 14.7, p<0.01 | 18.3, p<0.001 |
| Microstate syntax | Feature | A↔C/B↔D weakens | < 64 ch weakens | z=3.39–3.84, p<0.0025 | z=3.41–3.91, p<0.0025 |
| Effective connectivity | Feature | ↓ with density | < 64 ch degrades | 23.5, p<0.001 | 29.8, p<0.001 |
| Traveling waves | Feature | Speed underestimated | < 64 ch impaired | 16.8, p<0.01 | — |
| Functional hubs | Network | Hub set contracts | < 64 ch unreliable | J<0.5 at <64ch | J<0.5 at <64ch |

