## Supplementary Materials for "How Much Does Electrode Density Matter? A Multilevel Evaluation from EEG Signals to Cortical Networks"

| Channels | Density Label | Approximate System | Electrode names |
| --- | --- | --- | --- |
| 256 | High-density | 10–5 system | **Frontal (~60):** Fpz, AFz, Fz, AF1–AF8, F1–F8, AFF1–AFF8, FFC1–FFC8, plus dense intermediates.  **Central (~55):** Cz, FCz, CPz, C1–C6, FC1–FC6, CP1–CP6, FCC1–FCC6.  **Parietal (~50):** Pz, P1–P8, CP1–CP6, PPO1–PPO8, plus intermediates.  **Occipital (~45):** Oz, O1–O2, POz, PO1–PO8, I1–I2, plus dense posterior sites.  **Temporal (~46):** T7–T8, FT7–FT8, TP7–TP8, FTT7–FTT8, TPP7–TPP8, plus lateral dense sites |
| 128 | High-density | 10–10 system | **Frontal (~30):** Fpz, Fp1, Fp2, AF3–AF8, F1–F8, AFF1–AFF8, Fz **Central (~28):** FC1–FC6, C1–C6, Cz, CP1–CP6, CPz **Parietal (~26):** P1–P8, PPO1–PPO8, Pz, CP5–CP6 **Occipital (~20):** O1, O2, Oz, PO1–PO8, POz, I1–I2 **Temporal (~24):** T7, T8, FT7–FT8, TP7–TP8, FTT7–FTT8, TPP7–TPP8 |
| 64 | Medium-density | Extended 10–20 system | **Frontal (14):** Fpz, Fp1, Fp2, AF3, AF4, AF7, AF8, F1, F2, F3, F4, F7, F8, Fz  **Central (12):** FC1, FC2, FC3, FC4, C1, C2, C3, C4, Cz, CP1, CP2, CPz  **Parietal (12):** P1, P2, P3, P4, P5, P6, P7, P8, Pz, PPO1, PPO2, CP5  **Occipital (8):** O1, O2, Oz, PO3, PO4, PO7, PO8, POz  **Temporal (18):** T7, T8, FT7, FT8, TP7, TP8, plus lateral intermediates (FT9, FT10, TP9, TP10, etc.) |
| 32 | Low-density | Reduced 10–20 system | **Frontal (8):** Fpz, Fp1, Fp2, F3, F4, F7, F8, Fz  **Central (5):** C3, C4, Cz, FC1, FC2  **Parietal (4):** P3, P4, Pz, CPz  **Occipital (3):** O1, O2, Oz  **Temporal (12):** T7, T8, TP7, TP8, FT7, FT8, plus lateral intermediates (e.g., FC5, FC6, CP5, CP6, PO7, PO8) |
| 19 | Minimal-density | Standard 10–20 system | **Frontal (7):** Fp1, Fp2, F7, F3, Fz, F4, F8  **Central (3):** C3, Cz, C4  **Parietal (3):** P3, Pz, P4  **Occipital (2):** O1, O2  **Temporal (4):** T3 (T7), T4 (T8), T5 (P7), T6 (P8) |

***Table S1. Electrode configurations used for spatial subsampling (to be retained in the Supplementary Material).***

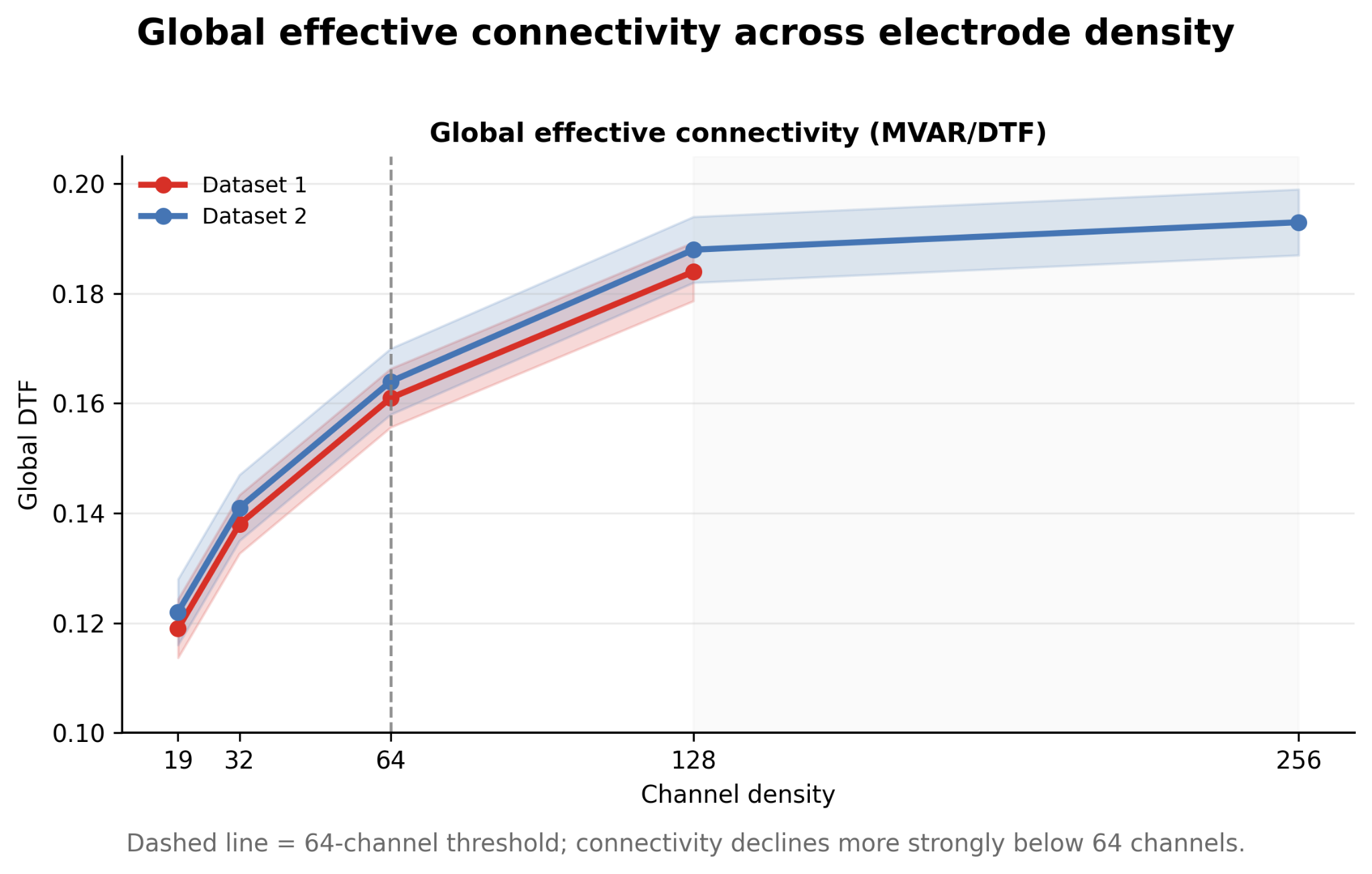

***Figure S1. Global effective connectivity across electrode density. Mean global MVAR/DTF values are shown for both datasets as channel count decreases from the highest-density configuration.***

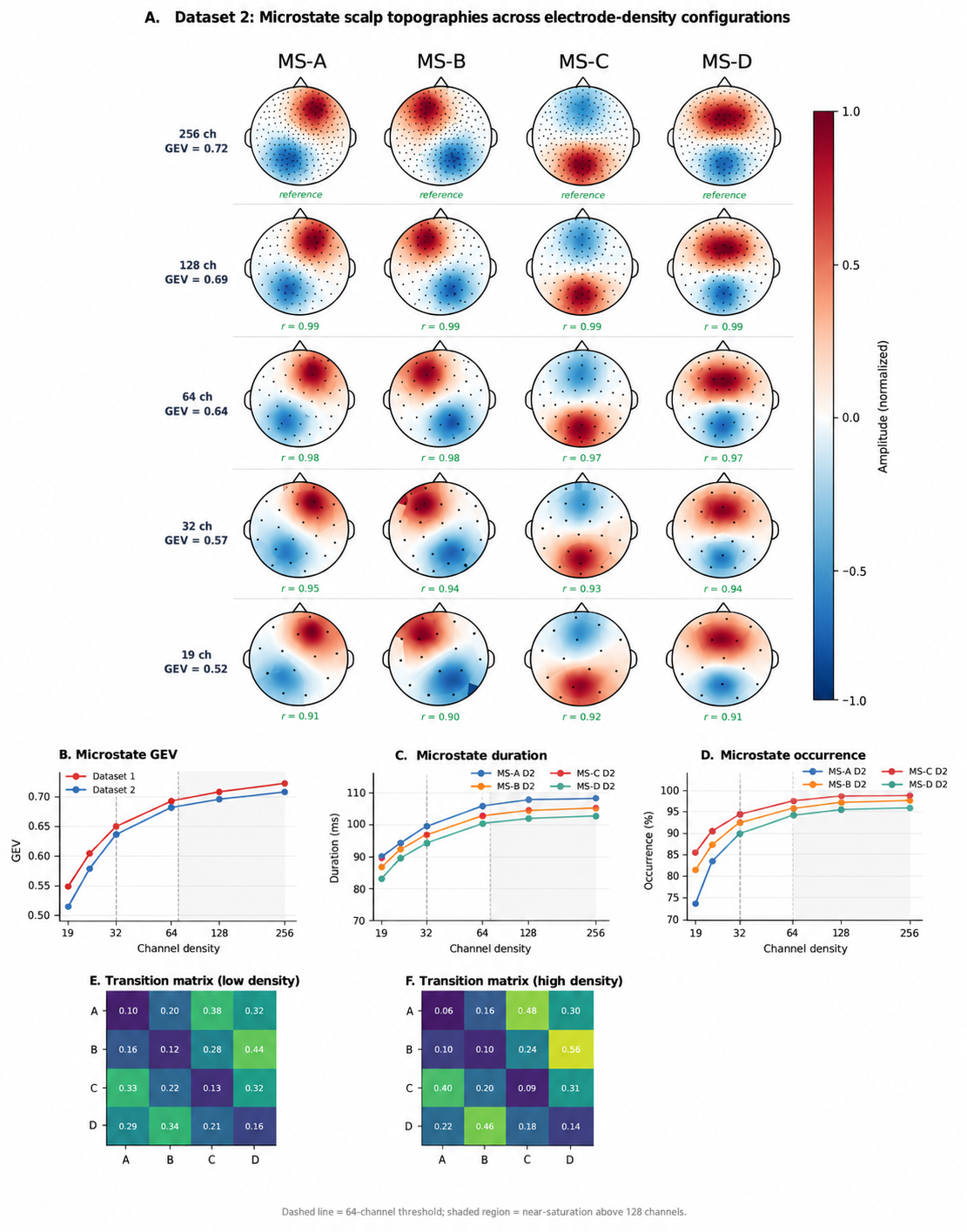

***Figure S2. Microstate topography and dynamics across electrode density. For Dataset 2, canonical microstate maps A–D are shown across 256-, 128-, 64-, 32-, and 19-channel configurations, together with density-dependent changes in global explained variance (GEV), duration, occurrence, and transition structure.***

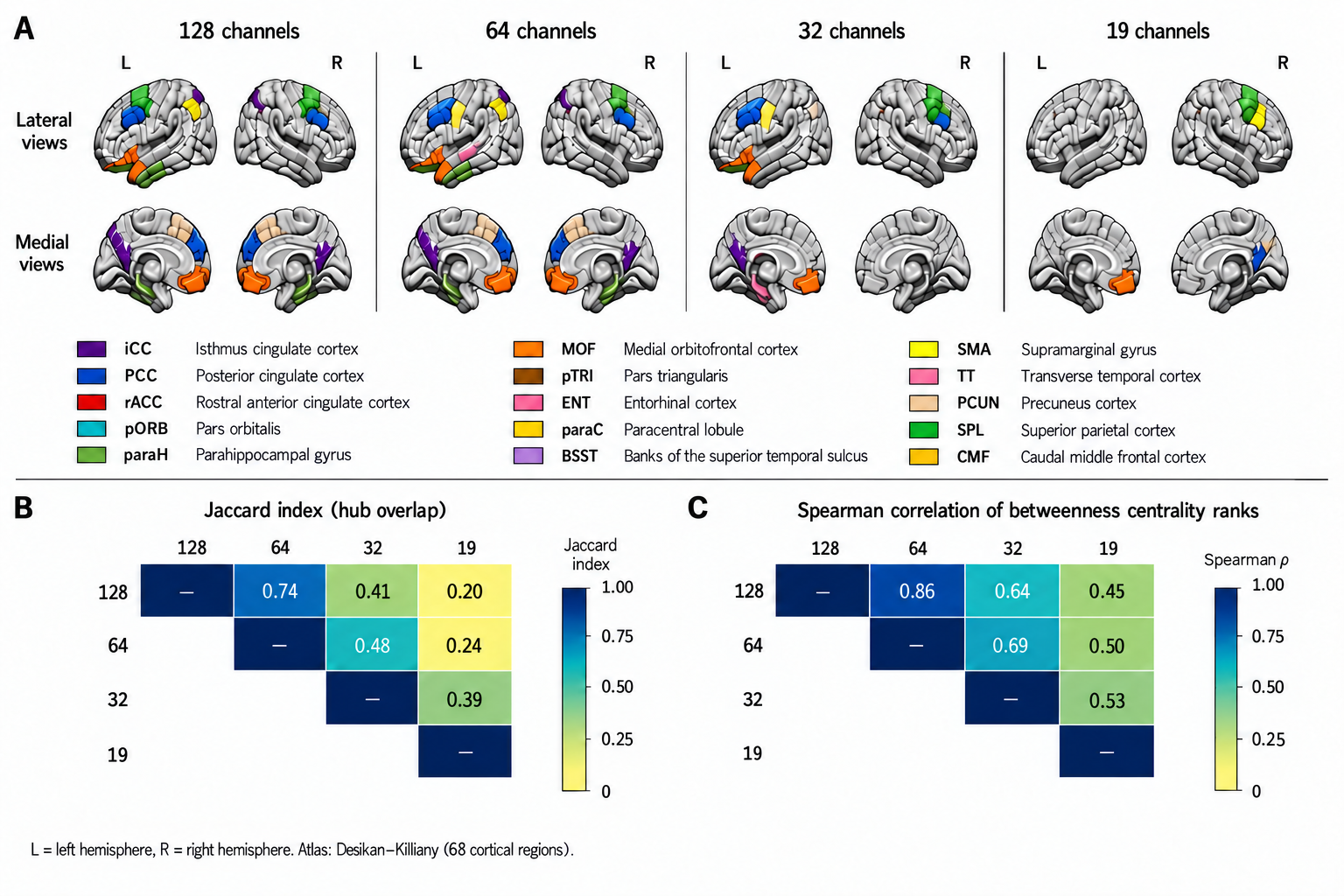

***Figure S3. Effect of electrode density on cortical hub identification and cross-density consistency in dataset 1. (A) Cortical regions identified as functional hubs across the 256-, 128-, 64-, 32-, and 19-channel configurations. Hubs were defined as regions belonging to the top 10% of betweenness-centrality values in source-level functional connectivity networks reconstructed on the 68-region Desikan–Killiany cortical atlas. Lateral and medial views of both hemispheres are shown, with colors indicating the anatomical identity of detected hubs. (B) Pairwise consistency of hub identification across electrode-density configurations. The Jaccard index quantifies overlap between the sets of detected hubs, whereas Spearman’s ρ quantifies preservation of the rank ordering of regional betweenness centrality across all cortical regions. L, left hemisphere; R, right hemisphere.***
